## Supplementary Code Listings & Results Tables for "TIAToolbox: An End-to-End Toolbox for Advanced Tissue Image Analytics"

### Supplementary Material

#### A. Code Listings

```python
156 from tiatoolbox.wsicore.wsireader import WSIReader
157 
158 tissue = WSIReader.open("path/wsi1.svs")
159 mask = tissue.tissue_mask()
160 
161 tissue_region, mask_region = (
162     wsi.read_rect(
163         location=(0, 0),
164         size=(512, 512),
165         resolution=0.5,
166         units="mpp",
167     )
168     for wsi in (tissue, mask)
169 )
```

*Listing A-1: A short Python expression demonstrating a synchronous reading of a tissue WSI and a lower resolution mask derived from the tissue WSI. Both read operations use the same function call arguments coordinates, including identical coordinates, as shown on lines 008 through 011. This is despite the underlying image data being stored at different resolutions. This demonstrates the power of the VirtualWSIReader object to enable easy reading from both a WSI and a derived image, such as a tissue mask, which may be internally represented at a different resolution.*

```python
from tiatoolbox.models.engine.patch_predictor import PatchPredictor

data = [img1, img2]  # input arrays as a list
predictor = PatchPredictor(
  pretrained model="resnet18-kather100k",
)
output = predictor.predict(
  data,
  mode="patch",
)
```

*Listing A-2: An example Python script for patch prediction using TIAToolbox using pre-extracted patches as input.*

```python
from tiatoolbox.models.engine.patch_predictor import PatchPredictor

predictor = PatchPredictor(
  pretrained model="resnet18-kather100k",
  pretrained_weights="path/weights.pth",
)
```

*Listing A-3: Code showing Python script of how to use your own pretrained weights, rather than those provided by TIAToolbox.*

```python
from tiatoolbox.models.engine.patch_predictor import PatchPredictor

# Input WSI file paths
data = ["path/wsi1.svs", "path/wsi2.svs"]

predictor = PatchPredictor(
  pretrained_model="resnet18-kather100k"
)

output = predictor.predict(
  data,
  mode="wsi",
)
```

*Listing A-4: An example Python script for patch prediction using TIAToolbox using whole-slide images as input. The same API is used for image tiles by changing the mode to "tile" in the predict method.*

```python
from tiatoolbox.models.engine.semantic_segmentor import SemanticSegmentor

# Input WSI file paths
data = ["path/wsi1.svs", "path/wsi2.svs"]

segmentor = SemanticSegmentor(
  pretrained_model="fcn_resnet50_unet-bcss",
)

# WSI prediction
output = segmentor.predict(
  imgs=[wsi_file_name],
  mode="wsi",
)
```

*Listing A-5: Supplying a WSI as input to a semantic segmentation model. Here, we use a U-Net with a ResNet50 backbone that is trained on the breast cancer semantic segmentation (BCSS) dataset.*

```python
from tiatoolbox.models.engine.nucleus_instance_segmentor import NucleusIns
tanceSegmentor

# Input WSI file paths
data = ["path/wsi1.svs", "path/wsi2.svs"]

# Instantiate the nucleus instance segmentor
inst_segmentor = NucleusInstanceSegmentor(
  pretrained_model="hovernet_fast-pannuke",
)

# WSI prediction
wsi_output = inst_segmentor.predict(
  [wsi_file_name],
  mode="wsi",
)
```

*Listing A-6: Supplying a WSI to a nucleus instance segmentation model. Here, we use a HoVer-Net trained on the PanNuke dataset.*

```python
from tiatoolbox.wsiscore.wsireader import WSIReader
from tiatoolbox.visualization.tileserver import TileServer

wsi = WSIReader.open("path/wsi1.svs")
app = TileServer(
  title="Testing TileServer",
  layers={
    "wsi1": wsi,
  },
)
app.run()
```

*Listing A-7: Creating a tile server web server gateway interface (WSGI) application and running a simple development HTTP server to display two WSI images in a web browser.*

```python
from shapely.geometry import Polygon
from tiatoolbox.annotation.storage import SQLiteStore, Annotation

# Store a simple triangle annotation
store = SQLiteStore("annotations.db")
annotation = Annotation(
  geometry=Polygon([(0, 0), (1, 0), (0, 1)]),
  properties={},
)
uuid = store.append(annotation)

# Access the annotation and add some properties
print(store[uuid])
annotation.properties = {"class": 1, "foo": "bar"}
store[uuid] = Annotation(triangle)

# Change just part of the properties
store.patch(uuid, {"class": 2})

# Query in a bounding box (left, right, top, bottom)
uuids = store.iquery([0, 0, 1, 1])

# Query with a predicate statement
results = store.query([0, 0, 1, 1], where="props['class']==2")
```

*Listing A-8: An example of creating an annotation and storing it in an SQLite database.  The annotation is given a universally unique identifier (UUID) as no key was specified. This is used to look up the annotation and modify it. Also shown here, is how to query for annotations using a bounding box and adding a predicate statement to filter the results.*

```python
import numpy as np

from tiatoolbox.models.engine.patch_predictor import PatchPredictor
from tiatoolbox.utils.misc import imwrite

WSI_PATH = "path/wsi1.svs"

# Tumour detection
tumour_predictor = PatchPredictor(
  pretrained_model='resnet18-idars-tumour',
)

tumour_output = tumour_predictor.predict(
  imgs=[WSI_PATH],
  mode='wsi',
)

tumour_mask = tumour_predictor.merge_predictions(
  WSI_PATH,
  tumour_output[0],
  resolution=5,
  units="power",
)
tumour_mask = tumour_mask == 2  # Binarise the output
imwrite('tumour_mask.png', tumour_mask.astype('uint8'))

# WSI Prediction
msi_predictor = PatchPredictor(
  pretrained_model='resnet34-idars-msi',
)

msi_output = msi_predictor.predict(
  imgs=[WSI_PATH],
  masks=['tumour_mask.png'],
  mode='wsi',
  return_probabilities=True,
)

# Slide-Level Score:
# Only consider MSI class
msi_probabilities = np.array(msi_output[0]['probabilities'])[...,1]
# Get the average over all tumour tiles
average_msi_probability = np.mean(msi_probabilities)
```

*Listing A-9: IDaRS inference using TIAToolbox. Here, we demonstrate that we can simplify the overall inference pipeline without the need for many lines of code. The following steps are performed: 1) tumor detection, 2) saving tumor mask, 3) mutation prediction in tumor regions and 4) patch aggregation.*

```python
import numpy as np
from matplotlib import pyplot as plt

from tiatoolbox.tools.graph import SlideGraphConstructor

# Load XY patch positions in the WSI
positions = np.load(f"path/wsi1.position.npy")
# Load features for each patch
features = np.load(f"path/wsi1.features.npy")

constructor = SlideGraphConstructor()
graph = constructor.build(
  positions[:, :2],
  features,
  feature_range_thresh=None,
)

constructor.visualise(graph)
plt.show()
```

*Listing A-10: Building and visualizing a slide graph from a set of patch locations and associated features.*

### Supplementary Material

#### B.Pretrained Models

*Supplementary Table A-1 Detailed metrics of Models for patch prediction.*

| Model Family | Architecture Variants | Training Dataset(s) | Metric(s) |
|---|---|---|---|
| **AlexNet**[1] | AlexNet | Kather 100k[2], PCam[3] | Supplementary Table A-5 |
| **ResNet**[4] | ResNet-18, ResNet-34, ResNet-50, ResNet-101 | Kather 100k[2], PCam[3] | Supplementary Table A-5 |
| **ResNeXt**[5] | ResNeXt-50, ResNeXt-101 | Kather 100k[2], PCam[3] | Supplementary Table A-5 |
| **Wide ResNet**[6] | Wide ResNet-50, Wide ResNet101 | Kather 100k[2], PCam[3] | Supplementary Table A-5 |
| **DenseNet**[7] | DenseNet121, DenseNet161, DenseNet169, DenseNet201 | Kather 100k[2], PCam[3] | Supplementary Table A-5 |
| **MobileNet**[8,9] | MobileNet-v2, MobileNet-v3 small, MobileNet-v3 large | Kather 100k[2], PCam[3] | Supplementary Table A-5 |
| **GoogLeNet**[10] | GoogLeNet | Kather 100k[2], PCam[3] | Supplementary Table A-5 |

*Supplementary Table A-2 Detailed metrics of Models for semantic segmentation.*

| Model Family | Architecture Variants | Training Dataset(s) | Metric(s) |
|---|---|---|---|
| **UNet**[11] | ResNet-50 backbone | TCGA-BCSS | Supplementary Table A-6 |
| **HoVer-Net**[12] | HoVer-Net+[13] | Private oral dysplasia cohort (not available) | Shephard *et al.*[13] |

*Supplementary Table A-3 Detailed metrics of Models for nucleus segmentation or classification.*

| Model Family | Architecture Variants | Training Dataset(s) | Metric(s) |
| --- | --- | --- | --- |
| **HoVer-Net**[12] | Original,<br>Fast,<br>HoVer-Net+ | Kumar (MoNuSeg Subset)[14],<br>PanNuke[15,16],<br>CoNSeP[12],<br>MoNuSAC[17],<br>Private oral dysplasia cohort (not available) | Graham *et al.*[12],<br>Gamper *et al.*[16],<br>Shephard *et al.*[13], |

*Supplementary Table A-4 Whole slide classification.*

| Model Family | Architecture Variants | Training Dataset(s) | Metric(s) |
| --- | --- | --- | --- |
| **IDaRS** | ResNet-18 (tumor)<br>ResNet-34 (mutation) | TCGA-COAD | Supplementary Table A-8 |
| **SlideGraph** | SlideGraph+ | TCGA-BRCA | Supplementary Table A-7,<br>Lu *et al.*[18] |

*Supplementary Table A-5: Patch classification performance of models provided by TIAToolbox on both the Kather100K and Patch Camelyon (PCam) datasets.*

| | Kather100K $F_1$ | PCam $F_1$ |
| --- | --- | --- |
| **AlexNet**[1] | 0.965 | 0.840 |
| **ResNet18**[4] | 0.990 | 0.888 |
| **ResNet34**[4] | 0.991 | 0.889 |
| **ResNet50**[4] | 0.989 | 0.892 |
| **ResNet101**[4] | 0.989 | 0.888 |
| **ResNeXt50 32x4d**[5] | 0.992 | 0.900 |
| **ResNeXt101 32x8d**[5] | 0.991 | 0.892 |
| **Wide ResNet50**[6] | 0.989 | 0.901 |
| **Wide ResNet101**[6] | 0.990 | 0.898 |
| **DenseNet121**[7] | 0.993 | 0.897 |
| **DenseNet161**[7] | 0.992 | 0.893 |
| **DenseNet169**[7] | 0.992 | 0.895 |
| **DenseNet201**[7] | 0.991 | 0.891 |
| **GoogLeNet**[10] | 0.990 | 0.899 |
| **MobileNet v2**[8,9] | 0.991 | 0.895 |
| **MobileNet v3 large**[8,9] | 0.992 | 0.890 |
| **MobileNet v3 small**[8,9] | 0.992 | 0.867 |

*Supplementary Table A-6: Semantic segmentation performance (Sørensen–Dice score) on the Breast Cancer Semantic Segmentation (BCSS) dataset. Here, we use a U-Net model with a ResNet50 encoder.*

| | **Tumor** | **Stroma** | **Inflammatory** | **Necrosis** | **Other** | **All** |
|---|---|---|---|---|---|---|
| **Amgad *et al.*[19]** | 0.851 | 0.800 | 0.712 | 0.723 | 0.666 | 0.750 |
| **TIAToolbox** | 0.885 | 0.825 | 0.761 | 0.765 | 0.581 | 0.763 |

*Supplementary Table A-7: Performance of SlideGraph+ using five-fold cross-validation provided as part of TIAToolbox, compared to the original implementation. Here we report the mean and the standard deviation across the folds.*

| | **HER2** | **ER** |
|---|---|---|
| **Lu *et al. (original)*** | 0.710±0.020 | - |
| **TIAToolbox** | 0.738±0.043 | 0.872±0.023 |

*Supplementary Table A-8: Performance of IDaRS provided as part of TIAToolbox, compared to the original implementation.*

| | **MSI** | **TP53** | **BRAF** | **CIMP** | **CIN** | **HM** |
|---|---|---|---|---|---|---|
| **Bilal *et al.*[20]** | 0.828 | 0.755 | 0.813 | 0.853 | 0.860 | 0.846 |
| **TIAToolbox** | 0.870 | 0.747 | 0.750 | 0.748 | 0.810 | 0.790 |
